# Multisite Test-Retest Reliability and Compatibility of Brain Metrics derived from FreeSurfer Versions 7.1, 6.0, and 5.3

**DOI:** 10.1101/2022.04.13.488251

**Authors:** Elizabeth Haddad, Fabrizio Pizzagalli, Alyssa H. Zhu, Ravi R. Bhatt, Tasfiya Islam, Iyad Ba Gari, Daniel Dixon, Sophia I. Thomopoulos, Paul M. Thompson, Neda Jahanshad

## Abstract

Automatic neuroimaging segmentation and parcellation tools provide convenient and systematic methods for extracting numerous features from brain MRI scans, and are becoming standard practice for large-scale coordinated studies. One such tool, FreeSurfer, provides an easy-to-use pipeline to extract metrics describing cortical and subcortical morphometry. Over the past two decades, there have been over 25 stable releases of FreeSurfer, and different versions are used across published works. Despite this, the reliability and compatibility of metrics derived from the most recent major version releases have yet to be assessed empirically. Here, we use test-retest data from three public brain MRI datasets to assess within-version reliability and between-version compatibility across 42 regional outputs from three versions of FreeSurfer: the latest, v7.1, and two previous stable releases - v5.3, and v6.0. We find v7.1 was less compatible with older versions for measuring cortical thickness. In particular, the thickness of the cingulate gyrus had low compatibility (intraclass correlation coefficient (ICC) between 0.37 and 0.61) between versions. Temporal and frontal poles, and the medial orbitofrontal surface area metrics, also showed low to moderate compatibility with v7.1. While our work compares all three versions, our sub-comparisons between the older versions (v5.3 and v6.0) replicates earlier findings of low compatibility of pallidum and putamen volumes. Low between-version compatibility was not always indicative of low within-version reliability – all versions showed good to excellent reliability across most regional measures (ICC>0.8). Age associations, quality control metrics, and Dice coefficients in an independent sample of 106 individual scans, processed with all three versions of FreeSurfer, revealed differences in results of downstream statistical analysis. As neuroimaging studies adopt more recently released software, we provide researchers with a reference to highlight the regions and metrics that may yield findings inconsistent with published works using older FreeSurfer software. An interactive viewer for the results is provided at http://data.brainescience.org/Freesurfer_Reliability/

## Introduction

The reproducibility of research findings in the biological sciences has recently come to light as a major problem, particularly for the neuroimaging-heavy fields of psychological and neuro-sciences (Button et al., 2013, Boekel et al., 2015, Bowring et al., 2019, Poldrack et al., 2020, Hodge et al., 2020). Studies on major depressive disorder (MDD), for example, have pointed out inconsistencies in results as well as difficulties in drawing comparisons due to analytical and study design variability (Stuhrmann et al., 2011, Dichter et al., 2015, Müller et al., 2017, Fonseka et al., 2018, Beijers et al., 2019, Kang et al., 2020). In one study, using a more heterogeneous sample and rigorous statistical testing, Dinga et al. (2019) were unable to replicate the statistical significance used to define MDD biotypes previously found in the literature. Inconsistent results investigating neuroimaging traits and diseases have also been found in studies of insomnia (Spiegelhalder et al., 2015) and mild traumatic brain injury (mTBI). A meta-analysis of 14 reports of working memory in mTBI showed mixed findings of functional magnetic resonance imaging (MRI) hyperactivity, hypoactivity, and some studies even report both hyper and hypo activity (Bryer et al., 2013). Neuroimaging offers mechanistic insights into the variability that leads to risk for brain dysfunction, yet these findings must be replicable in order to extend the use of MRI-derived biomarkers to a clinical setting.

It is important to understand how and why these discrepancies occur, so that we can better understand why certain findings are not reproducible. For example, studies may be underpowered, or the variable of interest might have different effects across populations. Experimental results can also be affected by methodological factors such as the type of data collection (Yan et al., 2020), data processing and analysis (Carp 2012, Bennett et al., 2013, Botvinik-Nezer et al., 2020, Lindquist 2020), tool version and selection (Gronenschild et al., 2012, Tustison et al., 2014, Dickie et al., 2017, Perlaki et al., 2017, Meijerman et al., 2018, Bigler et al., 2020, Zavaliangos-Petropulu et al., 2022), and even operating system environments (Glatard et al., 2015). The presence of pathological tissue has also been reported to cause systematic errors in segmentation output (Dadar et al., 2021). If sample population and methodology differ, it can be difficult to tease apart the main source of the discrepant findings.

Recent efforts in the neuroimaging community have heightened awareness and partially addressed concerns surrounding reproducibility. Guides and tools for enhancing reproducibility have been published in an effort to promote *Open Science*. Open science aims to provide transparency into research studies to better understand the data collected, the code implemented and software used, the analysis performed, and the full scope of results, including null findings (Zuo et al., 2014, Gorgolewski et al., 2015, Gorgolewski & Poldrack 2016, Poldrack et al., 2017, Nichols et al., 2017, Vicente-Saez & Martinez-Fuentes 2018, Kennedy et. al., 2019). These efforts often include detailed documentation and containerization of analytical software to ensure consistency of software version, and even operating system to the extent possible should the study be replicated. Large consortia, such as ENIGMA (Enhancing NeuroImaging Genetics through Meta-Analysis), have also addressed issues of low power and varying data processing pipelines by conducting large scale harmonized meta- and mega-analyses across international datasets (Thompson et al., 2020). Analytical protocols are proposed and approved by the community in advance; they are then distributed and made readily available. These protocols also include data quality control guidelines to improve analytic consistency across heterogeneous datasets and populations.

Large, publicly available and densely phenotyped datasets that use these protocols have recently become a powerful resource that have advanced the field of neuroscience (Horien et al 2021). Studies like the Alzheimer’s Disease Neuroimaging Initiative (ADNI) and the UK Biobank collect data from one thousand to tens of thousands of individuals (Weiner et al., 2015, Littlejohns et al., 2020) with some collecting longitudinal data that spans well over a decade (Weiner et al., 2017). Automatic segmentation tools are widely used on such datasets and have allowed for tens to hundreds of thousands of scans to be conveniently processed, thus enabling neuroimaging traits to be used in a wide range of clinical and epidemiological studies. However, these tools do not come without their challenges and limitations.

Data processed from updated versions of these softwares are continuously released (http://adni.loni.usc.edu/2021/) and this leaves researchers questioning which version is most reliable or whether data and results from work that used prior versions are compatible with those of later releases. If the detected effects depend on the software version used, then that variability could threaten the reproducibility of published research and compromise clinical translation. However, these version updates are often needed to keep up with the many advancements made in the neuroimaging field. For example, version updates may include added options or tools to work with higher resolution images, or more computational efficient image processing pipelines (e.g., the use of GPUs for processing). As newer software releases are made available, we often lack information on whether new results will be consistent with prior findings, and what the impact of a software upgrade will be. To understand sources of study variability, it is important to understand how version upgrades may impact outcome measures.

One such automatic feature extraction and quantification tool that is widely used in neuroimaging is FreeSurfer (Fischl, 2012). FreeSurfer is a structural MRI processing suite that allows researchers to obtain brain parcellations and metrics from just a single T1-weighted image. Running the software involves just a one command, but the process itself is quite extensive – where the single image undergoes 34 stepwise processing stages (https://surfer.nmr.mgh.harvard.edu/fswiki/recon-all). Notably, more than 60 research papers have been published detailing FreeSurfer’s algorithms and workflows (https://www.zotero.org/freesurfer/collections/F5C8FNX8). The overall processing steps include: image preprocessing, brain extraction, gray and white matter segmentation, reconstruction of the white matter and pial surfaces, labeling of cortical and subcortical regions, and a spherical nonlinear registration of the cortical surface using a stereotaxic atlas, allowing for a more accurate alignment of gyral and sulcal landmarks. Users can then extract features, such as cortical thickness (defined as the distance between the white matter and pial surfaces), surface area (or the area of all the triangles on the mesh representing the white matter surface), and cortical and subcortical volumes, measured in cubic millimeters (Fischl, 2012).

A PubMed search of “freesurfer”, in the year 2020 alone, results in a total of 344 publications, indicating its wide use as a neuroimaging resource (https://pubmed.ncbi.nlm.nih.gov/?term=%28freesurfer%29&filter=years.2020-2020). It has been a popular tool for over 20 years throughout which over 25 different stable releases have been disseminated (https://surfer.nmr.mgh.harvard.edu/fswiki/PreviousReleaseNotes). Version release updates have included, for example, improvements in accuracy of the cortical labels or a change/addition in a preprocessing step such as denoising or bias field correction (https://surfer.nmr.mgh.harvard.edu/fswiki/ReleaseNotes). These version changes may affect certain extracted measures. Gronenschild et al. (2012) compared volumes and cortical thickness measures across FreeSurfer v4.3.1, v4.5.0, and v5.0.0 and found many measurements differed significantly. After the release of the next version, v5.3, Dickie et al. (2017) performed correlation analysis between cortical thickness measures output from FreeSurfer v5.1 and v5.3, and found high compatibility between the two versions. Such work helped inform protocols for consortia such as ENIGMA, where groups that had run FreeSurfer versions older than v5.0, were asked to rerun their processing pipeline, whereas both v5.1 and v5.3 were used for analyses within certain working groups. A more recent study, Bigler et al. (2020), compared FreeSurfer v5.3 and v6.0 across a select set of volumes, finding low compatibility between versions within the globus pallidus.

The latest stable release, v7.1, has yet to be thoroughly assessed for intra-version reliability and between-version compatibility. Here, we assessed the reliability and compatibility of the last three stable FreeSurfer version releases – v5.3 (2013), v6.0 (2017), and v7.1 (2020) – across three publicly available test-retest datasets. We set out to determine the (1) between-version compatibility and (2) within-version reliability, for cortical thickness, surface area, and subcortical volumes. To test how these version differences may influence population-level findings, we ran all three FreeSurfer versions on a subset of cross-sectional data from the UK Biobank, a cohort of middle-aged to older adults. We visually quality controlled and computed Dice overlap scores between each pair of versions for all regional outputs. Finally, we determined the linear effect of age for each region and metric of interest, to understand the stability of this effect across software versions.

## Methods

### Datasets

Test-retest datasets from the Human Connectome Project (HCP) (Van Essen et al., 2013), Kennedy Krieger Institute (KKI) (Landman et al., 2010), and Open Access Series of Imaging Studies (OASIS-1) (Marcus et al., 2007) were used to assess reliability within and between FreeSurfer versions. We limited the analysis to 76 healthy individuals with T1-weighted brain MRI scans aged 19–61. KKI includes retest data from 21 healthy volunteers with no history of neurological conditions; a retest subset of 35 healthy young adults are provided by HCP, and OASIS-1 includes 20 nondemented subjects imaged twice. The max inter-scan interval of 11 months in the HCP dataset is longer than OASIS and KKI, yet we do not suspect considerable changes in brain structure between sessions given that HCP is comprised of young healthy adults between the ages of 22 to 35 (Van Essen et al., 2013).

See **Table 1** for more details.

**Table 1.** Cohort demographics and scan parameters for test-retest datasets analyzed. HCP is a family-based dataset including up to 4 individuals per family, so we limited our ICC investigations to one randomly chosen individual per family.

| Cohort | Age range; mean(SD) | No. of subjects (%F) | Max inter-scan interval in days (mean) | Manufacturer/Field strength | Voxel size [mm]<sup>3</sup> |
| --- | --- | --- | --- | --- | --- |
| HCP | 22–35; 30.7(2.97) | 35 (44%) | 330 (144) | Siemens 3T | [0.7 × 0.7 × 0.7] |
| KKI | 22–61; 31.8(9.47) | 21 (48%) | 14 | Philips 3T | [1 × 1 × 1.2] |
| OASIS | 19–34; 23.4(4.03) | 20 (60%) | 90 (20.6) | Siemens 1.5T | [1.0 × 1.0 × 1.25] |

A subset of 106 neurologically healthy individuals was selected from the UK Biobank (Miller et al., 2016) to test age association outcome differences between versions. This included 56 females with a mean age and standard deviation of 62.3 (7.2) years and 50 males with a mean age and standard deviation of 61.2 (7.7) years. In this case, being neurologically healthy was defined based on the following exclusion criteria: cancers and diseases of the nervous system, aortic valve diseases, head injuries, and schizophrenia/bipolar disorders. While the UK Biobank has over 40,000 individual scans, we selected a relatively small subset, with a sample size more in line with most single-site current neuroimaging studies.

## FreeSurfer Regions and Metrics of Interest

All scans were run through the same *recon-all* pipeline provided by FreeSurfer for stable v5.3, v6.0, and v7.1 releases. Cortical parcellations were computed based on the Desikan-Killiany atlas (Desikan et al., 2006), where 34 distinct regions on each cortical hemisphere are labeled according to the gyral patterns. For each cortical region, FreeSurfer outputs the average cortical thickness, surface area, and volume. We focus our analyses on cortical thickness and surface area, as these are largely independent measures (Winkler et al., 2010) and volume is a composite of the two. We also extract and evaluate the FreeSurfer derived measures of total intracranial volume (ICV) and volumes of eight subcortical regions: the nucleus accumbens, amygdala, caudate, hippocampus, lateral ventricle, pallidum, putamen, and the thalamus. These metrics are all ones that have been repeatedly used throughout multinational ENIGMA projects, and are therefore of particular interest to many collaborative investigators invested in reproducible findings. For all of our ICC analysis here, we report left and right measures, as well as average cortical thickness, total surface area, and average subcortical volumes. We also include hemisphere and whole brain cortical thickness and surface area.

## Statistics and Quality Control

Intra-class correlation coefficients (ICCs) were calculated using the *psych* library in R (https://CRAN.R-project.org/package=psych). The following three compatibility comparisons were evaluated: v7.1 vs. v6.0, v7.1 vs. v5.3, and v6.0 vs. v5.3. Only the first time points from the test-retest data were selected for these comparisons. ICC2 was used to compute between-version compatibility measures to account for any systematic errors using the following formula:

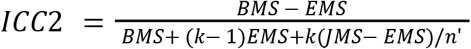

where *BMS* is the between-targets mean square, *EMS* is the residual mean square, *k* is the number of judges, *JMS* is the between-judges mean square, and n’ is the number of targets (in our context, the judges would correspond to different software versions used to compute the measures).

Within-version reliability measures were performed on within-subject test-retest data for FreeSurfer versions v7.1, v6.0, and v5.3. ICC3 was used to measure within-version reliability using the following formula:

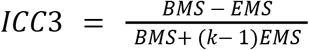

where *BMS* is the between-targets mean square, *EMS* is the residual mean square, and *k* is the number of judges. The reported ICC2 and ICC3 measures represent a weighted average to account for the number of participants in each dataset. ICC interpretation was based on Koo and Li (2016): ICCs<0.50 are considered poor; between 0.50 and 0.75 are moderate, between 0.75 and 0.90 denote good agreement; and values greater than 0.90 indicate excellent reliability.

To test if FreeSurfer version affects population level findings in studies of modest sample size, age associations were performed in a cross-sectional subset of the UK Biobank using linear regressions. Sex was used as a covariate; intracranial volume (ICV) was added as a covariate for subcortical volumes. In that same subset, detailed quality control (QC) was performed using the ENIGMA QC protocol (http://enigma.ini.usc.edu/protocols/imaging-protocols/) to test differences in regional fail rates across the versions. 54 subjects were assigned to rater #1 and 52 to rater #2. Each rater QC’ed the same subset across all three versions. Rater #3 then reviewed all QC fails for consistency. All subcortical QC was performed by rater #3 where a fail constitutes any notable over-or under-estimation of volume for any structure. Age associations were also performed in this QC’ed subset, where subjects were excluded if the QC of any ROI was inconsistent across versions. If subjects had consistent regional fails, they were kept in the analysis, but those regions were excluded. While many studies of such sample size may perform manual segmentation corrections, there is no way to ensure consistent manual editing across the outputs of all software versions. We therefore opted to exclude QC fails to ensure our reported differences were due to changes in software version.

For each set of regressions within a version, statistical significance was determined after controlling the false discovery rate at *q*=0.05 across 234 measures, which included all bilateral, unilateral, and full brain measures. False-discovery rate (FDR; Benjamini & Hochberg, 1995) corrected *p*-values and z-statistics were plotted on brain surfaces for comparison. All values, including uncorrected p-values, are tabulated on our web-viewer. Dice coefficients (Dice, 1945) were also calculated in the UK Biobank subset to assess the extent of spatial overlap of ROIs across versions, for all regions in the Desikan-Killiany atlas.

## Results

The full set of our reliability, compatibility, and association results are available through an interactive 3D brain viewer here: http://data.brainescience.org/Freesurfer_Reliability/.

### Between Version Compatibility

Version compatibility results between FreeSurfer v5.3, v6.0. and v7.1 for all cortical and subcortical metrics are shown in **Figure 1**. Overall, the version compatibility across all versions for average cortical thickness was good to excellent (ICC_v7.1:v6.0_=0.81; ICC_v7.1:v5.3_=0.85; ICC_v6.0:v5.3_=0.91). Similarly, left and right hemispheric thicknesses were good for v7.1 comparisons (left: ICC_v7.1:v6.0_=0.80, ICC_v7.1:v5.3_=0.86; right: ICC_v7.1:v6.0_=0.81, ICC_v7.1:v5.3_=0.83), and excellent when comparing v6.0 to v5.3 (left: ICC_v6.0:v5.3_=0.91; right: ICC_v6.0:v5.3_=0.90). Furthermore, version compatibility was excellent for v7.1 vs. v6.0 in several bilateral regional parcellations including the paracentral, postcentral, superior frontal, transverse temporal, and superior parietal cortices (ICC_v7.1:v6.0_>0.90). The postcentral (ICC_v7.1:v5.3_=0.91) and superior parietal (ICC_v7.1:v5.3_=0.91) gyri also showed excellent compatibility between v7.1 and v5.3. Additionally, v6.0 was highly compatible with v5.3 in the superior frontal, superior temporal, parahippocampal, supramarginal, *pars orbitalis*, and the banks of the superior temporal sulcus (ICC_v6.0:v5.3_≥0.90). Several bilateral regions showed poor compatibility between v7.1 and other versions, however. In particular, the lowest ICCs were found for the isthmus (ICC_v7.1:v5.3_=0.37; ICC_v7.1:v6.0_=0.58), posterior (ICC_v7.1:v5.3_=0.41; ICC_v7.1:v6.0_=0.55), caudal anterior (ICC_v7.1:v5.3_=0.46; ICC_v7.1:v6.0_=0.45), and rostral anterior (ICC_v7.1:v5.3_=0.61; ICC_v7.1:v6.0_=0.50) subregions of the cingulate gyrus. An example subject with notable differences in cingulate segmentations is displayed in **Figure 2A**. Other regions that showed moderate agreement with v7.1 and either v6.0 or v5.3 included the entorhinal (ICC_v7.1:v5.3_=0.64; ICC_v7.1:v6.0_=0.67), middle temporal (ICC_v7.1:v5.3_=0.68), and insular (ICC_v7.1:v6.0_=0.67) cortices, as well as the temporal (ICC_v7.1:v5.3_=0.69) and frontal poles (ICC_v7.1:v5.3_=0.70; **Figure 1A**).

**Figure 1.**
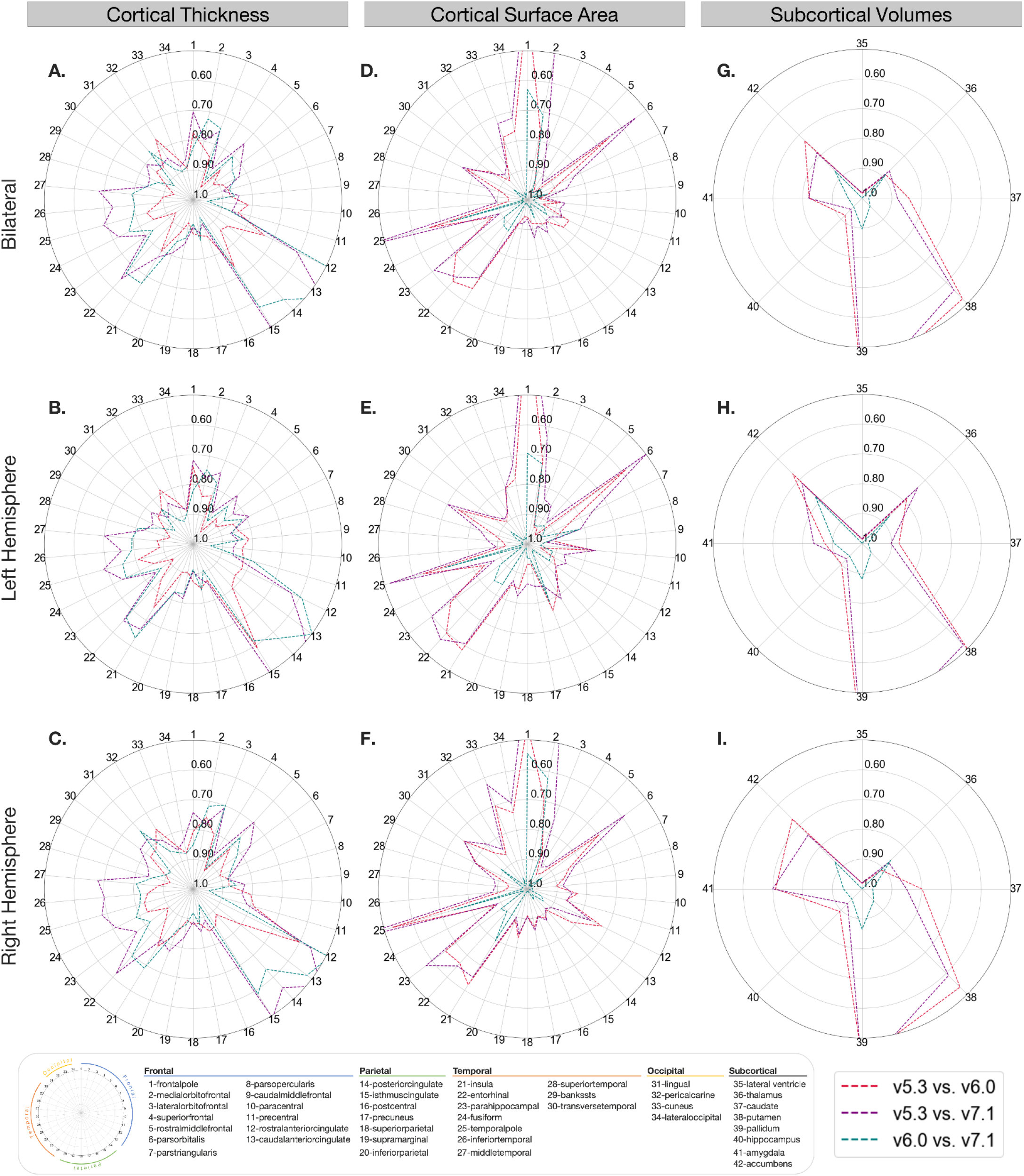
Regional compatibility ICC2 measures. Bilateral, left, and right ICC2 values comparing cortical thickness (**A, B, C**), cortical surface area (**D, E, F**), and subcortical volumes (**G, H, I**) between versions. Outer concentric circles represent lower ICC2 values, truncated at 0.50, while the center represents ICC2=1. Regions with the lowest compatibility differ for cortical thickness and surface area. These compatibility estimates shown are a sample-size weighted average of results in each of HCP, KKI, and OASIS datasets.

**Figure 2.**
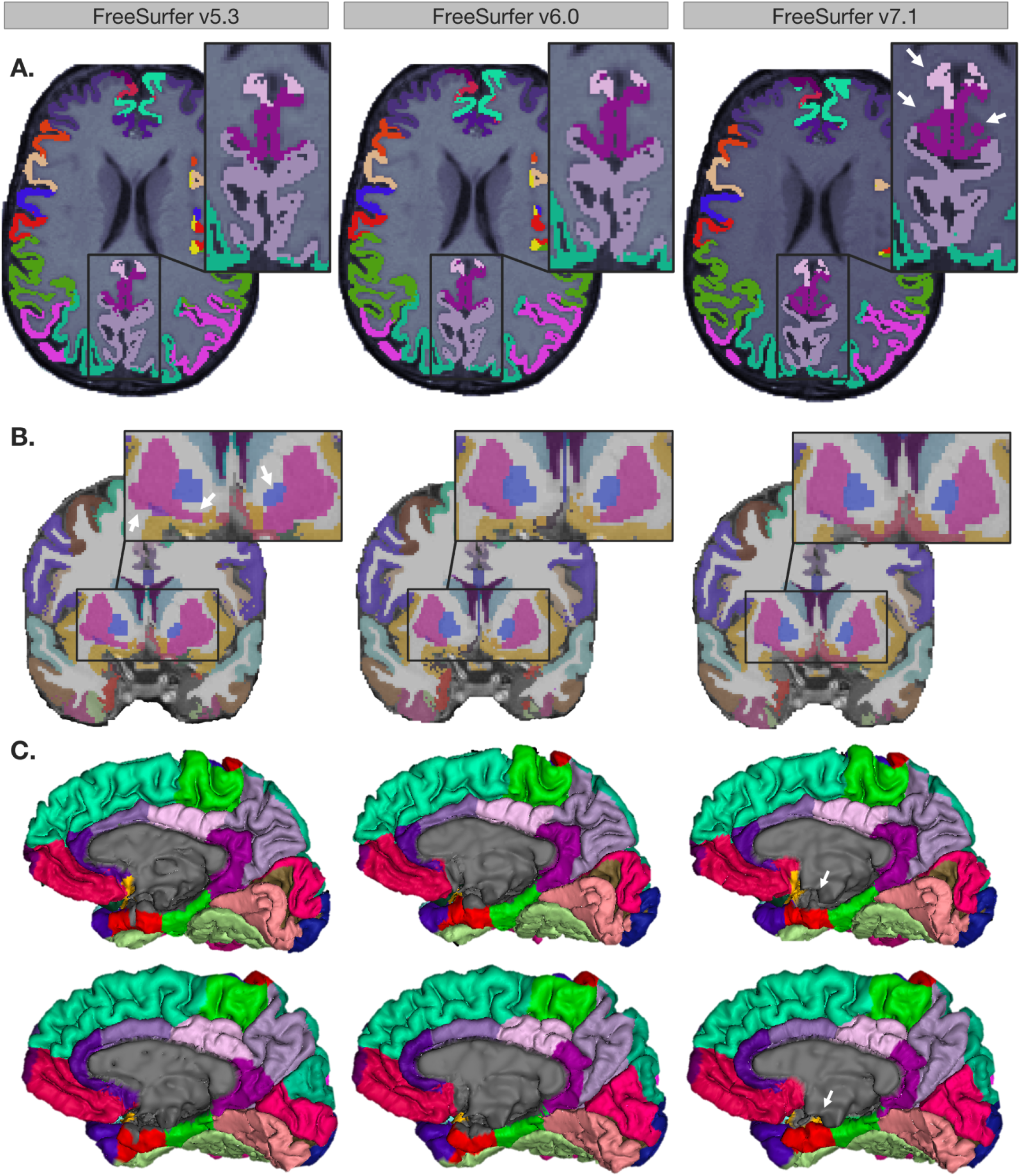
**A**. Axial slices from the same UK Biobank participant across versions. Arrows indicate posterior and isthmus cingulate differences in v7.1 vs. v5.3 and v6.0. **B**. Coronal slices from the same subject across versions. Arrows demonstrate v5.3 volume differences in the putamen and pallidum vs. v6.0 and v7.1. **C**. Medial surface representations of two UK Biobank participants across versions. Arrows highlight differences in the medial wall pinning, particularly in the entorhinal cortex, in v7.1 compared to the two prior releases.

Total surface area showed excellent compatibility across all three versions (ICC_v7.1:v6.0_=0.99; ICC_v7.1:v5.3_=0.96; ICC_v6.0:v5.3_=0.99). Left and right hemispheric surface area compatibility between versions were also excellent across all comparisons (ICCs>0.96). Overall, the two most compatible versions were v7.1 vs. v6.0, where, notably, 29/34 bilateral regions had ICCs>0.90. Several regions also showed excellent compatibility (ICC>0.90) across all three version comparisons: these included the caudal middle frontal, the inferior parietal, postcentral, posterior cingulate, rostral middle frontal, superior parietal, and the supramarginal gyri. However, we did find surface area compatibility discrepancies not only in regions mostly distinct from cortical thickness, but also between the pairs of versions being compared as well. The lowest bilateral regional surface area compatibility ICCs were observed in frontal and temporal areas when comparing newer versions to v5.3, where v7.1 showed lower compatibility to v5.3 than to v6.0. Frontal regions included the medial orbitofrontal cortex (ICC_v7.1:v5.3_=0.51; ICC_v6.0:v5.3_=0.76), *pars orbitalis* (ICC_v7.1:v5.3_=0.54; ICC_v6.0:v5.3_=0.66) and the frontal poles which were not compatible between either v7.1 (ICC_v7.1:v5.3_=0.19) or v6.0 (ICC_v6.0:v5.3_=0.32). However, compatibility between v7.1 and v6.0 was moderate for the medial orbitofrontal cortex (ICC_v7.1:v6.0_=0.71), excellent for the *pars orbitalis* (ICC_v7.1:v6.0_=0.94), and moderate for the frontal pole (ICC_v7.1:v6.0_=0.63). Temporal regions that followed similar trends included the parahippocampal gyrus (ICC_v7.1:v5.3_=0.61; ICC_v6.0:v5.3_=0.70) and the temporal poles (ICC_v7.1:v5.3_=0.43; ICC_v6.0:v5.3_=0.66), where in contrast v7.1 has excellent compatibility with v6.0 for the parahippocampal gyrus (ICC_v7.1:v6.0_=0.90) and moderate compatibility for the temporal pole (ICC_v7.1:v6.0_=0.73; **Figure 1D**).

ICV was highly compatible across all versions (ICCs>0.97). All bilateral subcortical volumes showed good to excellent compatibility when comparing v7.1 to v6.0 (ICCs>0.87). Good to excellent compatibility was also found comparing v5.3 to the newer versions in the lateral ventricle, hippocampus, thalamus, caudate, and amygdala (ICCs>0.82). Compatibility issues arose when comparing v7.1 and v6.0 against v5.3. Poor to moderate regional compatibility was found in the pallidum (ICC_v7.1:v5.3_=0.34; ICC_v6.0:v5.3_=0.36), putamen (ICC_v7.1:v5.3_=0.56; ICC_v6.0:v5.3_=0.52; **Figure 2B**), and to a lesser extent, the nucleus accumbens (ICC_v7.1:v5.3_=0.78; ICC_v6.0:v5.3_=0.73; **Figure 1G**).

### Within-Version Reliability

The meta-analyzed scan-rescan reliability for all cortical and subcortical metrics within each of FreeSurfer v5.3, v6.0. and v7.1 are shown in **Figure 3**. All versions showed high reliability for average bilateral, left hemispheric, and right hemispheric cortical thickness (ICC>0.90). Regional bilateral metrics with the lowest thickness ICCs – but still considered moderate to good – included the temporal pole (ICC_v7.1_=0.71; ICC_v6.0_=0.83; ICC_v5.3_=0.74), rostral anterior cingulate (ICC_v7.1_=0.83; ICC_v6.0_=0.79; ICC_v5.3_=0.78), and the medial orbitofrontal cortex (ICC_v7.1_=0.85; ICC_v6.0_=0.88; ICC_v5.3_=0.80; **Figure 3A**). Total bilateral, left hemispheric, and right hemispheric surface area reliability was also high (ICC=0.99) for all three FreeSurfer versions. The regions with the lowest surface area ICC were all still highly reliable, but included the frontal poles (ICC_v7.1_=0.88; ICC_v6.0_=0.87; ICC_v5.3_=0.77), insula (ICC_v7.1_=0.91; ICC_v6.0_=0.86; ICC_v5.3_=0.89), and entorhinal cortex (ICC_v7.1_=0.92; ICC_v6.0_=0.95; ICC_v5.3_=0.88) (**Figure 3D**). Regional bilateral subcortical volumes were all reliable for each of the three versions (ICC > 0.86; **Figure 3G**). ICV reliability was also very high (ICC>0.97) for all versions.

**Figure 3.**
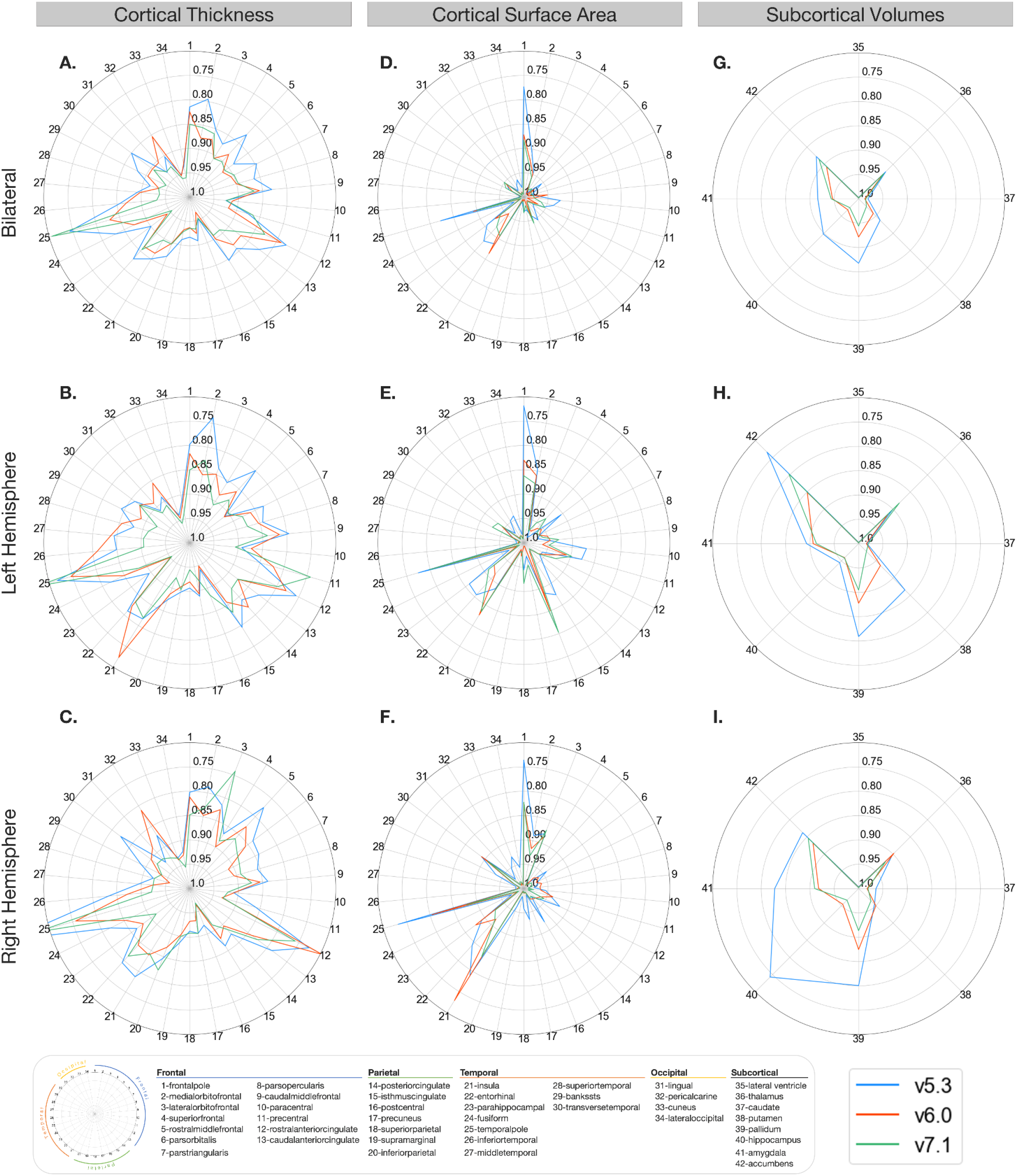
Regional reliability ICC3 measures. Bilateral, left, and right ICC3 values comparing cortical thickness (**A, B, C**), cortical surface area (**D, E, F**), and subcortical volumes (**G, H, I**) between versions. Outer concentric circles represent smaller ICC3 values, truncated at 0.70. Regions with the lowest reliability differ for cortical thickness and surface area. These reliability estimates shown are a sample-size weighted average of results in each of HCP, KKI, and OASIS datasets.

### Quality Control and Population-Level Analysis

Figure 4. highlights regional cortical quality issues noted in the subset of UK Biobank participant scans across each of the evaluated FreeSurfer versions. The region that showed the greatest difference in failure rate was the left superior temporal gyrus – where v7.1 performed the best (5.7% fails) followed by v6.0 (7.5% fails), and v5.3 performed the worst (12.3% fails; **Figure 4A**). In one subject with poor image quality, a general underestimation occurred throughout the brain in v5.3 but not in v6.0 and v7.1 (see **Figure 4B**). Other regions that failed at a relatively similar rate across all three versions included the left banks of the superior temporal sulcus (v7.1=17%; v6.0=17%; v5.3=18.9%), the left (v7.1=14.2%; v6.0=13.2%; v5.3=12.3%) and right (v7.1=11.3%; v6.0=12.3%; v5.3=13.2%) pericalcarine, the left middle temporal (v7.1=13.2%; v6.0=12.3%; v5.3=13.2%), the left cuneus (v7.1=13.2%; v6.0=12.3%; v5.3=10.4%), and the right cuneus (v7.1=8.5%; v6.0=10.4%; v5.3=10.4%).

The highest overlap was between v7.1 and v6.0 where most regions had a Dice coefficient of 0.90 or greater. The lowest overlap occurred when comparing v5.3 to both v7.1 and v6.0, particularly in the frontal pole (left: DC_v7.1:v5.3_=0.74, DC_v6.0:v5.3_=0.76; right: DC_v7.1:v5.3_=0.79, DC_v6.0:v5.3_=0.81), entorhinal (left: DC_v7.1:v5.3_=0.78, DC_v6.0:v5.3_=0.79; right: DC_v7.1:v5.3_=0.75, DC_v6.0:v5.3_=0.77), and right cuneus (DC_v7.1:v5.3_=0.75, DC_v6.0:v5.3_=0.77). Other regions with lower Dice coefficients were the cingulate regions, temporal pole, pericalcarine, and the banks of the superior temporal sulcus (**Figure 4C**).

**Figure 4.**
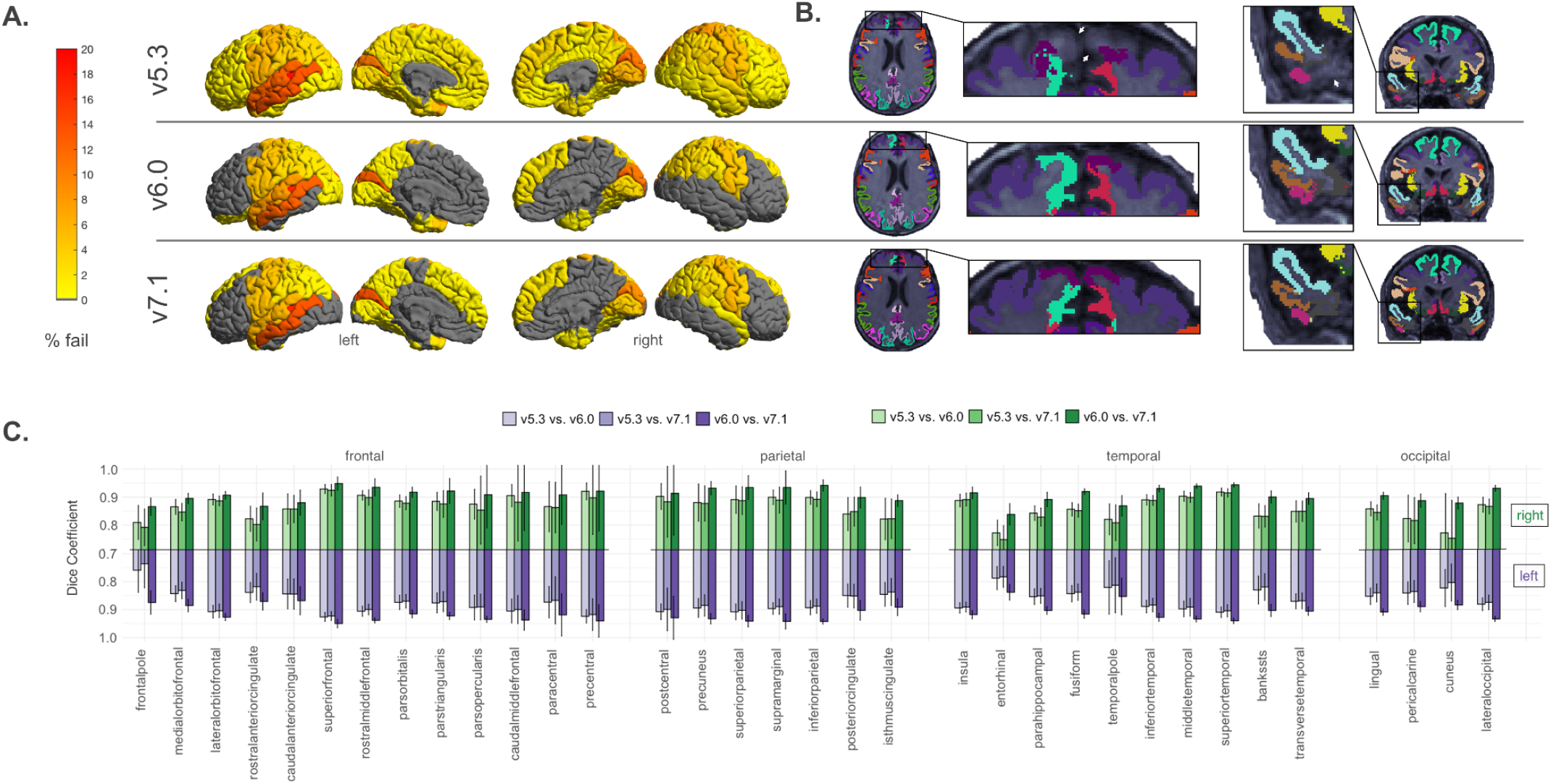
Cortical quality control results. Results based on 106 neurologically healthy UK Biobank participants. **A**. Manual cortical quality control results (percentage fail) based on the ENIGMA QC protocol across versions. Gray regions indicate no failures. Note more widespread failures particularly in the temporal and frontal regions due to a single subject in **B**. We also note generally higher rates of failure in the left temporal lobes across all versions. **C**. Dice scores across left and right hemisphere Desikan-Killiany atlas labels. We note the lowest overlap in the cuneus, entorhinal, pericalcarine, cingulate cortices, and temporal and frontal poles, particularly when comparing v5.3 to the newer versions.

Figure 5 highlights the subcortical quality issues noted across each of the evaluated FreeSurfer versions. The most regional failures were detected in v5.3. Failures occurred more often in the left hemisphere (**Figure 5A**). The most notable differences in failure rates were for the left pallidum (v7.1=0.9%; v6.0=0.9%; v5.3=18.9%), left amygdala (v7.1=7.5%; v6.0=11.3%; v5.3=17.9%), and left putamen (v7.1=0.9%; v6.0=1.9%; v5.3=14.2%). Example outputs may be viewed in **Figure 5B**. The regions with the lowest overlap were in the left and right pallidum (left: DC_v7.1:v5.3_=0.66, DC_v6.0:v5.3_=0.67; right: DC_v7.1:v5.3_=0.78, DC_v6.0:v5.3_=0.78) as well as the left and right nucleus accumbens (left: DC_v7.1:v5.3_=0.72, DC_v6.0:v5.3_=0.71; right: DC_v7.1:v5.3_=0.70, DC_v6.0:v5.3_=0.69) when comparing v5.3 to both newer versions. Notably, the segmentation of the left putamen often appeared larger and the left pallidum smaller in v5.3 compared to the newer versions (**Figure 5B**).

Age associations are shown in **Figure 6** and **Figure 7**. In the full set (106 UK Biobank scans) age associations for cortical thickness (**Figure 6**), v7.1 had 29 regions that survived FDR correction, less than both v6.0 with 32 and v5.3 with 43; all these regions showed lower thickness with age other than the right rostral anterior cingulate, which showed a positive association with age across all versions. The strongest associations were in the left supramarginal (*z*_v7.1_=-5.55, *q*_v7.1_=3×10^−5^; *z*_v6.0_=-6.24, *q*_v6.0_=1×10^−6^; *z*_v5.3_=-5.90, *q*_v5.3_=4×10^−6^) and left superior temporal (*z*_v7.1_=-4.88, *q*_v7.1_=1×10^−4^; *z*_v6.0_=-5.21, *q*_v6.0_=4×10^−5^; *z*_v5.3_=-5.33, *q*_v5.3_=2×10^−5^) for all three versions. All regions that were significant in v7.1 and v6.0 were also significant in v5.3, except for the left frontal pole in v6.0 (*z*_v7.1_=-2.19, *q*_v7.1_=9×10^−2^; *z*_v6.0_=-2.55, *q*_v6.0_=4×10^−2^; *z*_v5.3_=-1.83, *q*_v5.3_=1×10^−1^). Generally, v5.3 gave the lowest *p*-values and the highest effect sizes compared to v7.1 and v6.0. For the surface area age associations, no regions survived FDR correction in v7.1, whereas in v6.0 the left frontal pole survived correction, and in v5.3 the right paracentral, left banks of the superior temporal sulcus, right entorhinal, right lateral orbitofrontal, and right temporal pole were considered significantly associated with age after correction. For subcortical volumes, all regions were significantly associated with age, except for the left and right caudate and pallidum for all three versions and the right amygdala for v7.1 (*q*_v7.1_=0.08).

**Figure 5.**
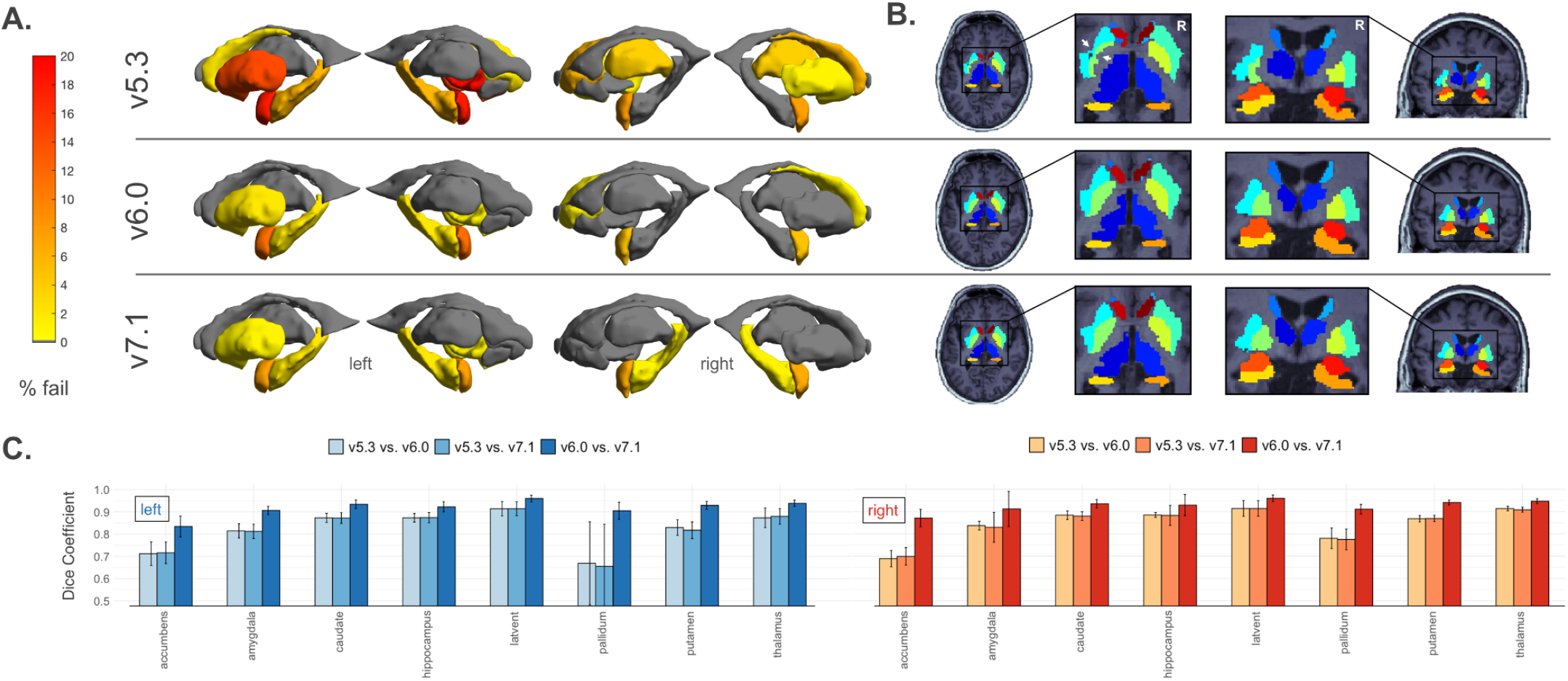
Subcortical quality control results. Results based on 106 neurologically healthy UK Biobank participants. **A**. Manual subcortical quality control results (percentage fail) across versions. Gray regions indicate no failures. Note generally higher fail rates in the left hemisphere and when comparing v5.3 to the newer versions. **B**. Example subcortical outputs. Arrows indicate the left putamen (cyan) and pallidum (light green) mis-segmentation in v5.3. **C**. Dice scores across left and right hemisphere subcortical regions. Note the lowest overlap when comparing v5.3 to v6.0 and v7.1.

**Figure 6.**
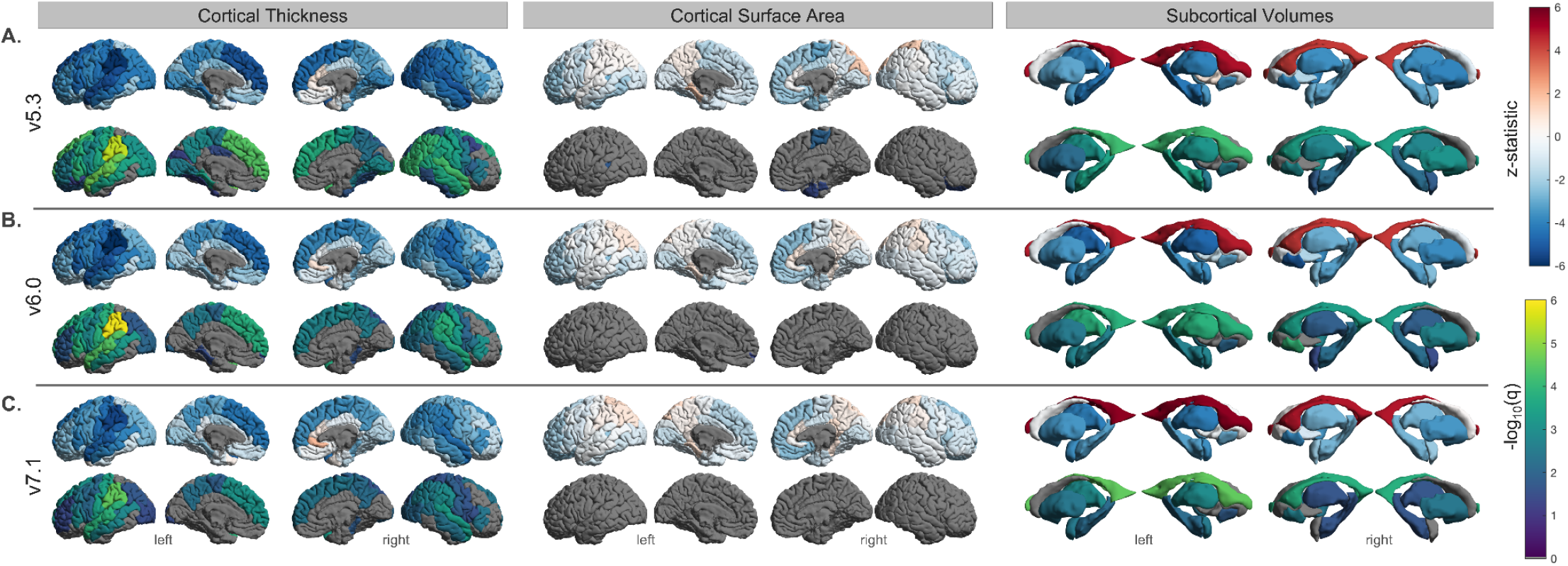
Regional age associations in all subjects. Results based on 106 neurologically healthy UK Biobank participants. **A**. FreeSurfer v5.3, **B**. v6.0, and **C**. v7.1. Top row indicates the effect sizes and bottom indicates -log_10_(*q*<0.05) for left and right surface area, thickness, and subcortical volumes. We note that v5.3 generally has the largest effect sizes, particularly for cortical thickness, and the largest number of statistically significant regions.

**Figure 7.**
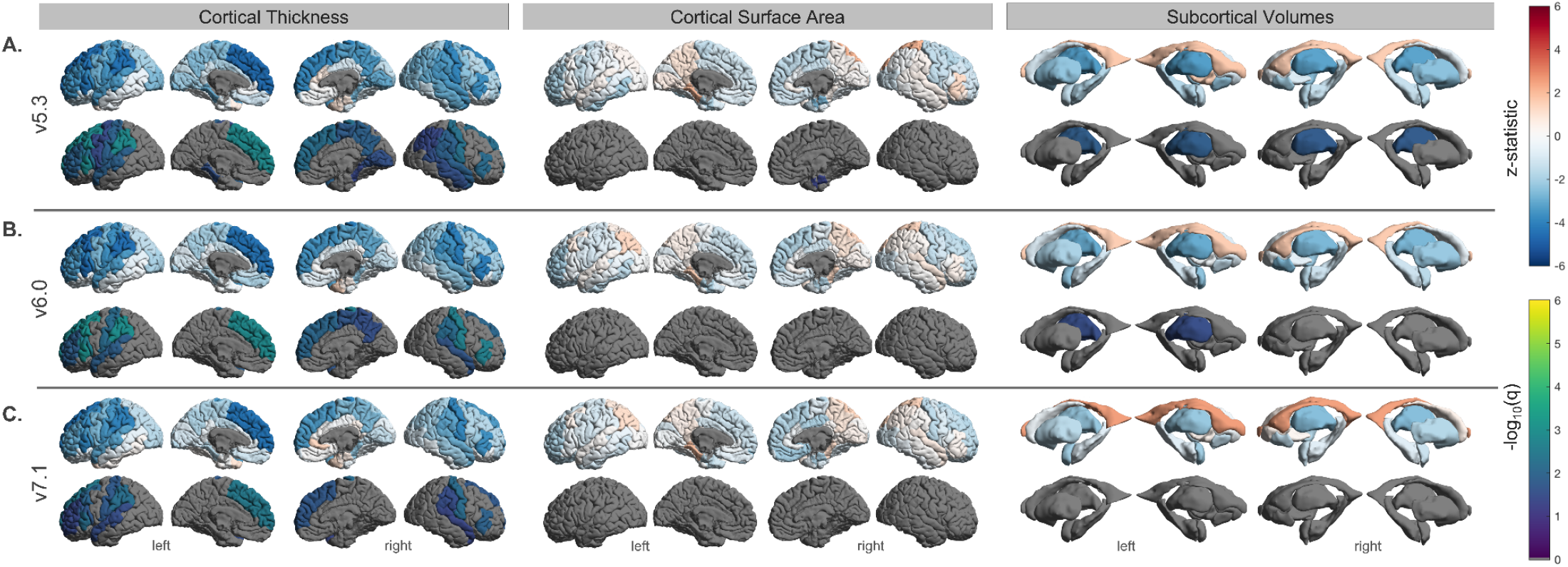
Regional age associations in subjects with no segmentation quality issues. Results based on n=69 (cortical) and 61 (subcortical) of the 106 neurologically healthy UK Biobank participants. **A**. FreeSurfer v5.3, **B**. v6.0, and **C**. v7.1. Top row indicates the effect sizes and bottom indicates -log_10_(*q*<0.05) for left and right surface area, thickness, and subcortical volumes. Several regions found to be significant in the full sample of n=106 did not survive FDR correction here.

A total of 69 subjects remained for regression analysis in the cortical QC’ed subset (37F, mean age: 61.1 ± 7.11). In this subset, cortical thickness was associated with age in 13 regions for v7.1, 16 for v6.0, and 22 for v5.3 (**Figure 7**). As with the full set above, all regions that survived FDR correction in v7.1 also survived in v6.0 and v5.3 and all regions that survived in v6.0 were also significant in v5.3. Cortical thickness regions that had a considerable proportion of fails and no longer reached the significance threshold in the QC’ed subset included the left banks of the superior temporal sulcus, left middle temporal, right precentral, and the right superior parietal gyrus. The left lingual, left cuneus, right pericalcarine, and the right banks of the superior temporal sulcus were all regions that had considerable quality issues and for which cortical thickness associations met FDR significance criteria for v5.3 in the full subset, yet these thickness associations were no longer significant in the QC’ed subset. The only surviving region in the QC’ed subset for surface area was the right entorhinal cortex in v5.3, although it is worth noting this region was not heavily QC’ed. The external surface in this area was apparently different in v7.1 compared to the previous versions (**Figure 2C**) and the rate at which this occurred would have resulted in the majority of participants being considered as a “fail” in the older versions.

61 subjects (35F, mean age: 62.6 ± 6.8 years) were found to have no quality issues in the subcortical segmentations across any versions. Age associations with these subjects indicated that only the thalamic volumes were significantly associated with age in v5.3 (both right and left) and v6.0 (left only).

## Discussion

Our work has four main findings that may help explain how a FreeSurfer version upgrade may impact results:

1. The compatibility between v7.1 and the previous version, v6.0, was largely good to excellent for measures of cortical surface area and subcortical volume, with the exception of the medial orbitofrontal cortex and the frontal/temporal poles. Most compatibility issues arose in regional cortical thickness estimates, where moderate or even poor compatibility was seen in the thickness estimates of the cingulate gyrus (rostral anterior, caudal anterior, posterior, and isthmus), entorhinal, insula, and orbitofrontal regions (medial and lateral).
2. There were substantial compatibility issues between v7.1 and v5.3, in cortical regional thickness, area, and subcortical volume. Thickness measures with low compatibility between v7.1 and v5.3 were the same as those between v7.1 and v6.0, suggesting these thickness compatibility issues have been more recently introduced with the latest 7.1 version. Regions with cortical surface area and subcortical volume compatibility issues between v7.1 and v5.3 were the same as the regions that were less compatible between v5.3 and v6.0, suggesting these area and volume differences were introduced with v6.0, not v7.1.
3. The test-retest reliability for all v7.1 metrics evaluated here was good to excellent, except for the temporal pole. This is in line with the reliability previously established in v6.0 and v5.3, and again confirmed here.
4. Age associations revealed generally smaller effect sizes in v7.1 compared to earlier releases, where v5.3 detected the largest effects overall. Quality issues were more prevalent in v5.3, particularly in the left superior temporal gyrus, pallidum, and putamen. Age associations did not meet the statistical significance threshold in many of the heavily quality controlled regions.

The regions in which v7.1 had the lowest compatibility with the previous versions were along the caudal-rostral axis of cingulate cortex. The subdivisions of the cingulate cortex play distinct roles in large-scale brain networks including the visceromotor, ventral salience, dorsal executive/salience, and default mode networks (Touroutoglou & Dickerson 2019). Alterations in the subregions of the cingulate cortex have been demonstrated throughout the lifespan and in association with different neuropsychiatric disorders. For example, compared to controls, developmental delays in adolescents with attention deficit hyperactivity disorder are seen most prominently in the thickness of the prefrontal regions including the cingulate cortices (Vogt 2019). In post traumatic stress disorder (PTSD) studies, the anterior midcingulate, and in some cases the posterior cingulate, show, on average, lower thickness in individuals with PTSD compared to healthy controls (Hinojosa et al. 2019). Subregions of the cingulate cortex have also been associated with age related cognitive performance. In “SuperAgers”, or adults over the age of 80 years, whose episodic memory is resistant to age-related decline, a preservation of the anterior cingulate thickness is observed (Harrison et al., 2012, Gefen et al., 2015, Sun et al., 2016, Harrison et al., 2018, de Godoy et al., 2021). Many of these studies were performed using versions of FreeSurfer that precede v7.1, so possible replication issues in future studies may be partially explained by the version incompatibility described in this work.

Other regions with lower thickness compatibility with v7.1 included the medial and lateral orbitofrontal, entorhinal, and insular cortices. Inferior frontal regions such as the medial and lateral orbitofrontal cortices are often susceptible to signal loss and bias field inhomogeneities. v7.1 uses an updated bias field and denoising method that could affect the gray/white matter contrast in these areas. Temporal regions, such as the entorhinal and insular cortex, which were less compatible with v7.1, could be due to an algorithmic update that pins the pial surface in the medial wall to the white matter surface. This prevents a premature cutoff through the hippocampus and amygdala, which may affect surrounding regions in earlier versions. Notably, visual inspection of the external surface of the entorhinal cortex revealed an improvement of the entorhinal pinning to the medial wall in v7.1 – as opposed to prior versions (**Figure 2C**). This issue was extremely prevalent, and considering these subjects as “QC-fail” would have resulted in the majority of subjects failing; therefore subject scans affected by cutoff in v5.3 and v6.0 remained included in our “error-free” subset. Downstream effects of this may be demonstrated in our age associations within the full n=106 sample. Here, the left insular thickness showed significant age effects in v5.3 and v6.0, as well as the thickness of the right entorhinal cortex in v5.3, but neither showed associations with age in v7.1. The entorhinal cortex plays an important role in mediating information transfer between the hippocampus and the rest of the brain (Garcia & Buffalo 2020, Coutureau & Scala 2009). Measurements of its thickness are widely assessed in Alzheimer’s disease, as it is one of the first regions to be impacted in the disease (Braak & Braak 1991) and researchers have found associations between its thickness and markers of amyloid and tau (Thaker et al., 2017). Entorhinal thickness is often a feature of interest in models that are designed to predict progressive cognitive decline due to its early vulnerability and role in the prodromal stages of Alzheimer’s disease. Although v7.1 may have a more anatomically accurate segmentation, we advise caution when comparing the performance of predictive models that use earlier releases of FreeSurfer for deriving this metric.

Compatibility issues between v7.1 and older versions were less frequent with surface area and did not occur in the same regions as cortical thickness. This could be due to the relative independence of these measures: surface area is calculated as the area of all the triangles on the white matter surface, the large area covered by many surfaces makes them more robust to slight variation in vertex counts. On the other hand, cortical thickness is measured as the distance between the vertices of the white matter and pial triangulated surfaces, and is often between 2-4mm thick, a span of only 2-4 voxels; slight variability in partial voluming may have a more dramatic effect on cortical thickness, yet as the thickness is averaged in the entire area, a slight variation in the number of vertices on the surface will have little effect on the averaged cortical thickness estimates. The independence of these measures has also been established in relation to their genetic associations (Winkler et al., 2010, Grasby & Jahanshad et al., 2020) overall suggesting that our results are not unexpected. Measures of v7.1 surface area that had poor compatibility with v5.3 (and moderate with v6.0) included the frontal and temporal poles. The release of v7.1 included a remeshing of the white matter surface to improve triangle quality in the surface mesh models – potentially impacting the most rounded points of the frontal and temporal lobes. We find that v7.1 had the lowest fail rate in the temporal pole compared to v5.3 and v6.0 suggesting an improvement in the parcellation.

Subcortical volumes are also another set of metrics derived from FreeSurfer that are of major interest to neuroimaging researchers (Satizabal et al., 2019, Ohi et al., 2020). Efforts to provide references of normative subcortical volume changes that occur as a result of aging have been put forth (Potvin et al., 2016, Coupé et al., 2017, Narvacan et al., 2017, Dima et al., 2022, Miletić et al., 2022, Bethlehem et al., 2022). For example, Potvin et al., 2016 pooled data from 21 research groups (n=2790) and segmented subcortical volumes using FreeSurfer v5.3 to provide norms of volumetric estimate changes during healthy aging. Although this study, along with many others, provides a valuable resource to researchers, we advise caution with the newer versions when referencing normative data derived from v5.3, particularly in the lentiform nucleus. The lentiform nucleus (i.e., the putamen and globus pallidus combined) has often been found to be difficult to segment due to the high white matter content in the pallidum – making it more difficult to distinguish gray-white matter contrast (Ochs et al., 2015, Visser et al., 2016, Makowski et al., 2018, Bigler et al., 2020). We find poor compatibility in the pallidum and moderate in the putamen when comparing v7.1 and v5.3. Visual QC of these regions revealed a higher failure rate and lower Dice overlap in v5.3 compared to v7.1, particularly in the left hemisphere. However, we find the compatibility between v7.1 and v6.0 to be excellent and the Dice overlap was greater than 90% in the lentiform nucleus. This suggests that changes made in the release of v6.0 contributed to v5.3 discrepancies. For example, the putamen does not extend so far laterally in the two newer versions — a known issue noted in the release notes of v6.0.

The main goal of our work was to evaluate FreeSurfer’s latest stable release, v7.1, yet it is also worth noting how v6.0 differs from v5.3. While compatibility was generally good for cortical thickness, regional surface area estimates were more moderately compatible, with the frontal pole even showing poor compatibility, similar to v7.1 compared to v5.3. Temporal lobe regions showing moderate compatibility in surface area between v6.0 and v5.3 included the entorhinal, insula, parahippocampal, and temporal pole. Updates that accompanied the release of v6.0 that may contribute to these compatibility discrepancies include improved accuracy of the cortical labels and an updated template (*fsaverage*) that “fixes” the peri/entorhinal labels. As previously mentioned, v6.0 compatibility with v5.3 was poorest in the pallidum and putamen. Our results coincide with Bigler et al. (2020) where the lowest agreement was also found in the pallidum and putamen when comparing v5.3 to v6.0.

One limitation of our study was that there was no available higher-resolution or post-mortem ground truth data to know which FreeSurfer version most represents true anatomical structure. However, given that many of these measures have been widely studied regarding their relationship with age, even in the absence of postmortem or higher resolution data (Salat et al., 2004, Fischl, 2012, Frangou et al., 2022), we instead assess age associations to gauge the downstream consequences of version differences. Here, QC of regional parcellations was performed to rule out any spurious associations with gross missegmentations. One example worth noting is that v7.1 and v6.0 may be better able to handle images with lower quality and/or motion as evidenced by one subject in our UK biobank subset that failed in v5.3 but not with the newer versions (**Figure 4B**). This could be due to the improved error handling of the Talairach registration: if one registration fails, v7.1 and v6.0 would try an older atlas. Another example involved the left middle temporal gyrus, which is often susceptible to underestimations due to the spillage/overestimation of the banks of the superior temporal sulcus into that gyrus. This occurred at approximately the same rate across versions. When associating the thickness of both the left banks of the superior temporal sulcus and the middle temporal gyrus with age before quality control, all versions reveal significant associations for both regions. After removing subjects encountering this issue, although the direction of the effects stayed the same, neither region was associated with age in any of the versions. While this may be due to a reduced sample size and study power, it is also possible that findings in these regions may not represent true anatomical structure, and may instead be due to common segmentation errors. It is also worth noting that our results are solely based on the Desikan-Killiany (DK) atlas (Desikan et al., 2006) and translation to other atlases may not apply. We chose the DK atlas as it consists of a set of coarse regions defined by anatomical landmarks that can be reasonably quality controlled. Most other atlases, while possibly more precise, define finer parcellations based on cortical function, connectivity, topography, myelin, or a combination thereof (Glasser et al., 2016, Schaefer et al., 2018). Visual quality control by region may not be readily possible when cortical parcellations are finer and there are over 100 regions in each hemisphere, so version performance of segmentation accuracy may be more difficult to compare. Our datasets were exclusively from adults without major neurological abnormalities, so our findings may not necessarily generalize to cohorts of young children, adolescents, or individuals with significant brain abnormalities.

Overall, we find generally high within-version reliability across most versions, and many advantages to using FreeSurfer v7.1 over older versions for adult neuroimaging studies. However, considerable differences are observed when analyzing between-version compatibility for regional cortical thickness, surface area, and subcortical volumes. It is important to consider these compatibility differences when pooling data or statistical inferences across software versions, and when comparing findings across published works, especially for those regions with lower compatibility. Understanding these differences may help researchers to make informed decisions on study design and provide insight into reproducibility issues.

## Acknowledgments

This research was funded by NIH grants R01MH117601 and R01AG059874. HCP data were provided [in part] by the Human Connectome Project, WU-Minn Consortium (Principal Investigators: David van Essen and Kamil Ugurbil; U54MH091657) funded by the 16 NIH Institutes and Centers that support the NIH Blueprint for Neuroscience Research and by the McDonnell Center for Systems Neuroscience at Washington University. KKI was supported by NIH grants NCRR P41 RR015241 (Peter C.M. van Zijl), R01NS056307 (Jerry Prince), R21NS064534 (Bennett A. Landman/Jerry L. Prince), R03EB01246 (Bennett A. Landman). OASIS: Cross-Sectional: Principal Investigators: D. Marcus, R. Buckner, J. Csernansky, J. Morris; P50 AG05681, P01 AG03991, P01 AG026276, R01 AG021910, P20 MH071616, and U24 RR021382.

